# TC10 differently controls the dynamics of Exo70 in growth cones of cortical and hippocampal neurons

**DOI:** 10.1101/2024.06.03.597209

**Authors:** Hiteshika Gosain, Karin B. Busch

**Author notes:** **Statement of significance** Exo70 is a central subunit of the vesicle tethering complex in exocytosis. Exocytosis is important, for example, in the regeneration of nerves. The Rho-GTPase TC10 is a regulator of exocytosis, but how TC10 influences Exo70 is not fully understood. Here, we used single-molecule tracking to quantify the effect of TC10 on the dynamics of Exo70 at the plasma membrane. We showed that TC10 slowed Exo70 dynamics in the axonal growth cone of cortical neurons consistent with stimulated vesicle binding, but found no obvious effect of TC10 on Exo70 dynamics in hippocampal neurons. Moreover, the effect of TC10 on Exo70 was again different in non-polar HeLa cells, suggesting that the control of Exo70 by TC10 is a complex process.

## Abstract

The exocyst is an octameric protein complex that acts as a tether for GOLGI-derived vesicles at the plasma membrane during exocytosis. It is involved in membrane expansion during axonal outgrowth. Exo70 is a major subunit of the exocyst complex and is controlled by TC10, a Rho family GTPase. How TC10 affects the dynamics of Exo70 at the plasma membrane is not well understood. There is also evidence that TC10 controls Exo70 dynamics differently in nonpolar cells and axons. To address this, we used super-resolution microscopy to study the spatially resolved effects of TC10 on Exo70 dynamics in HeLa cells and the growth cone of cortical and hippocampal neurons. We generated single particle localization and trajectory maps and extracted mean square displacements, diffusion coefficients, and alpha coefficients to characterize Exo70 diffusion. We found that the diffusivity of Exo70 was different in nonpolar cells and in the growth cone of neurons. TC10 stimulated the mobility of Exo70 in HeLa cells, but decreased the diffusion of Exo70 in the growth cone of cortical neurons. In contrast to cortical neurons, TC10 overexpression did not affect the mobility of Exo70 in the axonal growth cone of hippocampal neurons. These data suggest that mainly exocyst tethering in cortical neurons was under the control of TC10.

## Introduction

Exocytosis is a dynamic process contributing to cellular autophagy, polarization, membrane expansion and vesicle fusion. During early neuronal differentiation, neurite outgrowth and extension are key events **[1, 2]**, that require membrane surface expansion by exocytosis at the leading edge **[3]**. The tethering of exocytic vesicles is mediated by the exocyst complex, which is composed of two subcomplexes: the Exo70 subcomplex with Sec10, Sec15 and Exo84, and a subcomplex composed of Sec3, Sec5, Sec6 and Sec8 **[4]**. Exo70 is the determinant for the assembly of the whole complex and thus determines exocytosis. Dynamics of Exo70 at the PM is mediated by TC10, a member of the Rho family of GTPase proteins **[5]**. GTP hydrolysis of TC10 promotes neurite outgrowth by promoting exocytosis **[6, 7]**. GTP hydrolysis of TC10 promotes tethering of vesicles and subsequently the release of Exo70 from the plasma membrane. It is not clear, whether the effect of TC10 on Exo70 dynamics is quantitively the same in all neurons, since other Rho GTPase also modulate axon formation **[8]**. To address this, we compared the effect of TC10 on Exo70 dynamics in cortical and hippocampal neurons. We analyzed the dynamics of Exo70 at the plasma membrane in in early-stage differentiating neurons using super resolution imaging and quantitative analysis. We quantified the mobility of Exo70 in the presence of TC10 and mutants of TC10 by single particle (SPT) tracking using TIRF microscopy. The constitutively active TC10/Q67L (or TC10CA) and the fast-cycling TC10/34L (TC10FC) were used to study the influence of mutants on Exo70 dynamics. Our study revealed that TC10-mediated control of Exo70 dynamics is distinct between hippocampal and cortical neurons.

## Material and methods

### Cell culture and cultivation

HeLa cells (purchased from DSMZ, # ACC 57) were chosen as a nonpolar model cell line. HeLa cells were cultured in Dulbecco’s Modified Eagle Medium (DMEM), 10% Fetal Bovine Serum (FBS), 1% non-essential amino acids (NEA), 1% HEPES buffer solution, 1% L-Glutamine and 1% Penicillin as an antibiotic. Table 1 lists general material used and Table 2 culture media composition.

**Table 1:**
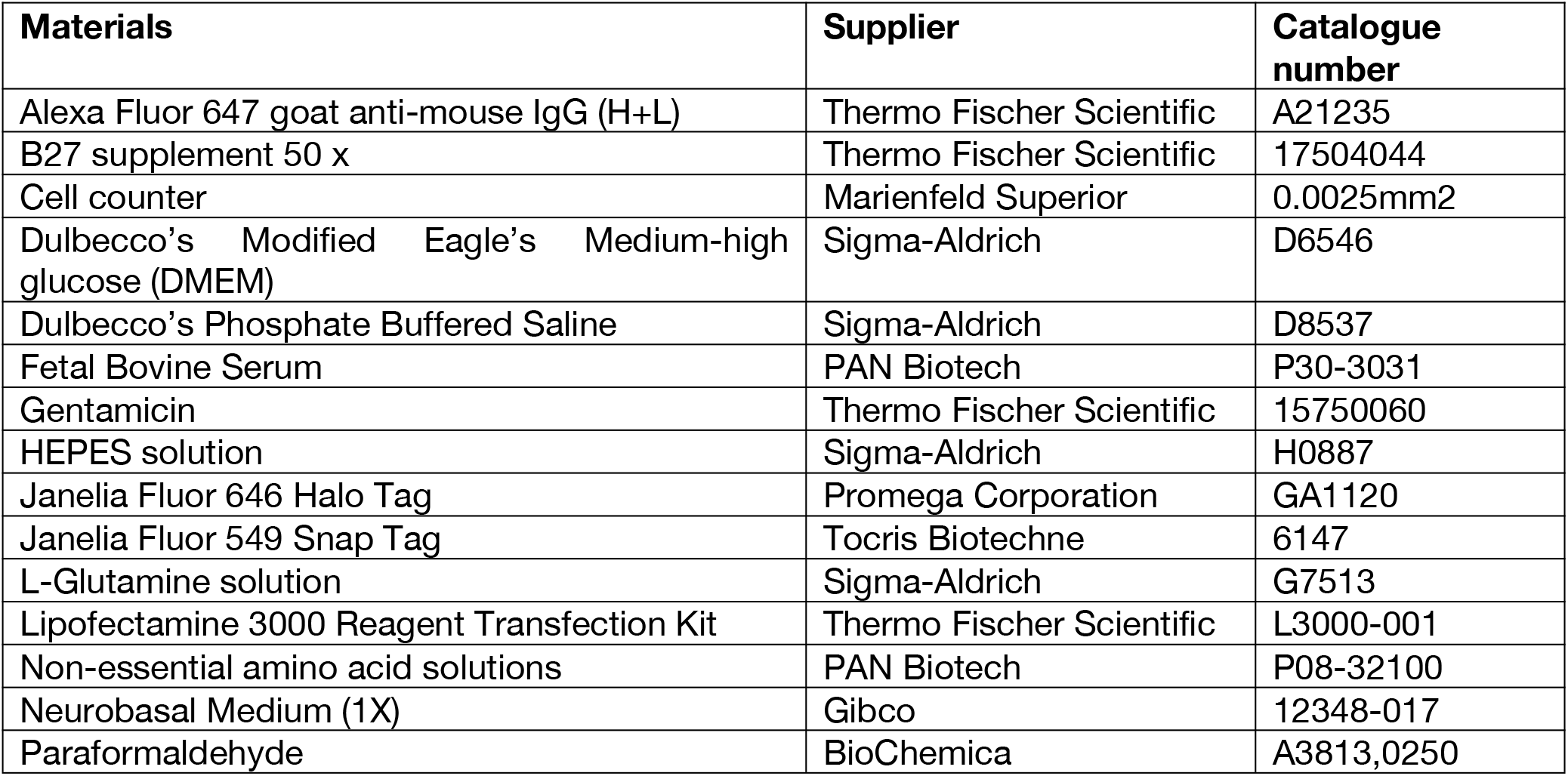
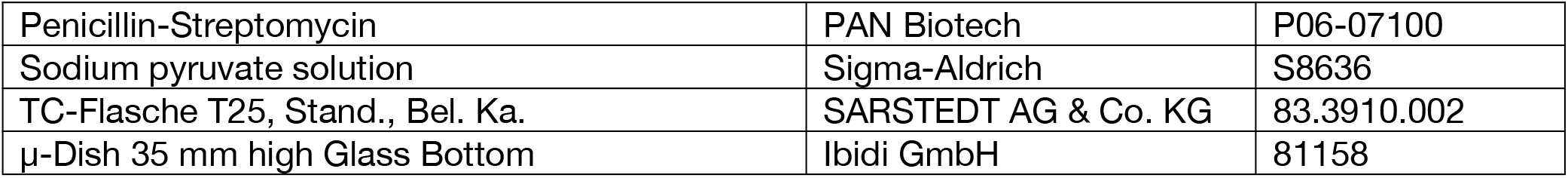
General Material.

**Table 2:** Cell culture media composition.

| Media | Composition |
| --- | --- |
| DMEM++ (growth medium) | 1x DMEM with Phenol red, 10% heat-inactivated FBS, 2 mM L-Glutamine, NEAA (1%), HEPES (1 μM) and 50 μg/mL Penicillin streptomycin |
| DMEM+++ (imaging medium) | 1x DMEM without Phenol red, 10% heat-inactivated FBS, 2 mM L-Glutamine, NEAA (1%), HEPES (1 μM) and 50 μg/mL Penicillin streptomycin |
| NBM++ (conditioned medium) | 50 mL Neurobasal medium (NBM), 1 mL B27, 500 μl 100x Glutamine and 125 μl Gentamicin |

**Table 3:** List of antibodies for Immunofluorescence staining.

| Antibody target | specification | Provider | Ordering number |
| --- | --- | --- | --- |
| Exo70 | Mouse monoclonal | Santa Cruz | sc-365825 |
| TC10 | Rabbit polyclonal | Proteintech | 17805-1-AP |
| Alexa Fluor 488 | goat anti-rabbit | Thermo Fischer Scientific | A-11008 |
| Alexa Fluor 555 | goat anti-mouse | Thermo Fischer Scientific | A21147 |

Primary rat embryo neurons were obtained from a rat Hippocampus on embryonic day 18 (E18). All animal protocols were performed in accordance with the guidelines of the North Rhine-Westphalia State Environment Agency (LANUV). Rats were maintained at the animal facility of the Institute for Integrative Cell Biology and Physiology (University of Münster) under standard housing conditions with a 12 h light/dark cycle at constant temperature (23°C) and with ad libitum supply of food and water. Timed pregnant rats were set up in house. Pregnant rats were anesthetized by exposure to isoflurane followed by decapitation and primary cultures prepared from embryos at embryonic day 18 (E18). During dissection, neurons from all embryos (regardless of sex) were pooled.

Neurons were grown in supplemented Neurobasal medium (NBM++) and kept at 37°C and 5% CO_2_. To obtain polar neurons, Hippocampi were isolated from rat embryos (at day 18 of embryonic development) and collected in HBSS on ice. Hippocampi were incubated at 37°C for 8 minutes in 2 mL Trypsin (not re-suspended). They were washed five times in pre-warmed DMEM++ before being resuspended in 2 mL DMEM++ with a 1 mL pipette tip after aspirating trypsin. Neurons were then seeded on poly-ornithine coated dishes with NBM++ medium and incubated at 37°C with 5% CO_2_.

For microscopy, 100,000 neurons were seeded on poly-ornithine coated 3 cm glass bottom dishes. Neurons were cultured in DMEM++ medium, which was replaced with conditioned media after 3 hours and placed in an incubator at 37 °C and 5% CO_2_. The day when neurons were seeded is termed as “Day in vitro zero” (DIV0). The first day following seeding is known as DIV1, and the days after that are known as DIV2, DIV3, and so on.

### Transfection

Hela cells were transfected with Lipofectamine 3000 according to the manufacturer’s instructions, neurons by the calcium phosphate transfection procedure. After washing, neurons were supplied with pre-collected conditioned medium and left for three days to mature. At DIV3, live neurons were imaged.

### Staining self-labeling tags for live cell imaging

The protein of interest (POI) was genetically fused to either the HaloTag (HaLo) or the Snaptag (SNAP) at the C-terminus of the Exo70 protein sequence resulting in Exo70-HaLo and the N-terminus of TC10 (SNAP-TC10). For HaloTag-labeling, we used a fluorescent derivative of 1-chlorohexane as a substrate, the HaloTag ligand (HTL), which forms a covalent bond with the HaloTag. For Snaptag-labeling, a fluorescent benzyl-guanine (BG) was used. Janelia Fluor® (JF) dyes JF646-HTL (500 pM) and JF549-BG (2 nM) were used in single molecular studies performed with the TIRF microscope. Cells were incubated with the dye-substrates for 30 minutes at 37°C and 5% CO_2,_ washed and imaged in pre-warmed imaging medium. For confocal imaging, 50 nM JF646-HTL and 50 nM JF549-BG were used.

### Immunostaining

Cells were fixed with 4% paraformaldehyde (PFA) containing 15% sucrose for 10 minutes at room-temperature (RT). Samples were washed three times in PBS, then treated with 50 mM Ammonium Chloride (NH_4_Cl) to eliminate free aldehyde groups. Cells were permeabilized with 0.1% TritonX-100 in PBS for 10 min. After washing with PBS, blocking was done with 2% goat serum and 3% bovine serum albumin (BSA) (1 hour at RT in a dark, humid environment). Residual blocking buffer was aspirated after 1 hour. The primary antibody against endogenous Exo70 (12014-1-AP, Proteintech) was used at a dilution of 1: 50. The secondary antibody conjugated with Alexa Fluor 555 or Alex Fluor 488 was used at a dilution of 1: 500.

### Cloning of Exo70-HaLo

Exo70 was amplified from pEGFP-C3-Exo70 (Addgene plasmid #53761) using the primers 5’-CGGAATTCTG ATGATTCCCC CACAGGAGG-3’ and 5’-GCGTCGACAC GGCAGAGGTG TCGAAAAGGC-3’ and was inserted into the pSEMS-HaloTag-vector .

### Microscopy

For confocal imaging, an inverted cLSM (Leica TCS SP8 FLIM system) was used. The microscope was equipped with a unique Acoustic Optical Deflector (AOD), two photomultiplier tubes (PMT), two highly sensitive hybrid GAsP (HyD) detectors, a 63x water objective (NA 1.2) and a pulsed white light laser (WLL). Measurements were done at 37°C and 5% CO_2_. An image format of 1024×1024 pixels was used with a scan speed of 700 Hz to obtain images with high signal to noise ratio. z-stacks were acquired and processed in FIJI to create an averaged intensity projection of all the images in the stack.

### TIRF microscopy

An inverted TIRF microscope (Xplore, Olympus) was used for SPT studies. The microscope is equipped with a TIRF objective (100× oil, NA of 1.49), an Optosplit IV (Cairn) for simultaneous recording of four channels and four diode lasers (wavelengths of 405 nm, 488 nm, 560 nm, and 640 nm) connected by an optical fiber to four independently controlled TIRF modules. To reduce wide-field illumination, a collimated beam of light was generated by focusing into the back focal plane of the objective. The output intensities of the diode laser in the system were controlled on a microsecond timescale by an acoustical optical tunable filer (AOTF). To image single molecules, we used a highly sensitive sCMOS camera (ORCA Fusion– BT, Hamamatsu) at a pixel size of 65 nm. 5,000 frames with a constant time interval (58.8 Hz) were usually recorded. The image format for a single channel was set to 500x500 pixels.

### Single particle tracking and data processing

For tracking analysis, the raw images were processed in FIJI using the Trackmate plugin to extract track information for the analysis ^26^. This program takes the simplest approach possible by detecting each particle’s nearest neighbors in a circular region in successive frames. Spots were detected using the “LoG detector” filter. In combination with the median filter and subpixel localization, the threshold was modified according to the signal intensity. The data was visualized using the “HyperStack Displayer” in uniform color. The tracking was done using the simple LAP tracker (linking conditions: maximum distance, gap-closing conditions, maximum distance). The generated tracks were further filtered using a “Number of spots in tracks” filter. This filter was used to make sure the tracks were long to reduce statistical errors and discard tracks produced by blinking occurrences. Resulting tracks (saved as a “.XML” file) were further analyzed in Matlab using the @msdanalyser to get ensemble MSD curves, diffusion coefficient distributions, and alpha coefficient distributions ^33^. Generally, ensemble MSD curves were plotted after weighted averaging over all particles. Diffusion coefficients were extracted from the MSD curves using power law and fitted with a linear function. The first 25% (200 ms) of the trajectory were fitted. The threshold for the quality of the fit were R2 coefficients > 0.8. The fits gave us the values of and for individual tracks according to the models described by Qian ^34^.

### Statistical analysis

For statistical analysis, Origin™ software was used for analysis of *a* and 𝒟 Usually, the Kruskal-Wallis ANOVA test was followed by post-hoc analysis.

## Results

First, we checked whether Exo70 and TC10 colocalize. Therefore, we immuno-stained both proteins in HeLa cells. TC10 and Exo70 were found in the plasma membrane (PM) but also in puncta-like structures in the cytosol. About 45% of Exo70 and TC10 signals colocalized indicating interaction (Figure S1). To test, whether TC10 affects Exo70 dynamics, we expressed tagged versions of Exo70 and TC10 in HeLa cells and cortical neurons. This allowed us to conduct single particle localization (SPL) and tracking (SPT) to resolve the effect of TC10 on the spatiotemporal behavior of Exo70 at the plasma membrane under TIRF excitation. For live cell imaging, Exo70 was genetically fused to the self-labeling Halo-Tag at its C-terminus annotated as Exo70-Halo. TC10 was genetically fused to the SNAP-Tag at N-terminus, called as SNAP-TC10. Halo-Tagged and SNAP-tagged proteins are self-labeling constructs and are commonly used for super-resolution microscopy [9]. SNAP-TC10 was labeled with JF-549-Snap, Exo70-HaLo with JF-646-Halo.

### TC10 overexpression results in increased mobility of Exo70 in non-polar cells

First, we determined the impact of TC10 on Exo70 mobility in non-polar HeLa cells. Using Total Reflection Fluorescence Microscopy (TIRFM), we excited Exo70 exclusively at the plasma membrane (PM). We recorded 5000 frames with a frame rate of 59 Hz, localized the molecules and generated trajectories (**Figure 1A-B**). From these, Exo70 mobility was determined. In addition to wildtype TC10, two TC10 mutants with different activity based on the GDP/GTP state were generated using point mutations. The TC10 point mutated at Q67L mutant was permanently in the GTP binding state and thus constitutively active (TC10CA) **[10]**. The mutant TC10F42L rapidly switches between GDP and GTP states and thus is fast cycling (TC10FC) **[11]**. Mean square displacements (MSD) were calculated for each trajectory. We compared ensemble time averaged MSDs for Exo70 in the presence of TC10 and its variants in the short time diffusion. As typical for particles at the PM, longer recording shows confinement (**Figure 1C**). Calculation of MSD vs. time on a logarithmic scale shows the deviation from normal behavior (**Figure 1D**). To determine the diffusion coefficient *D* (-log D; µm^2^/s), log MSDs were fitted with a linear function as given by

**Figure 1:**
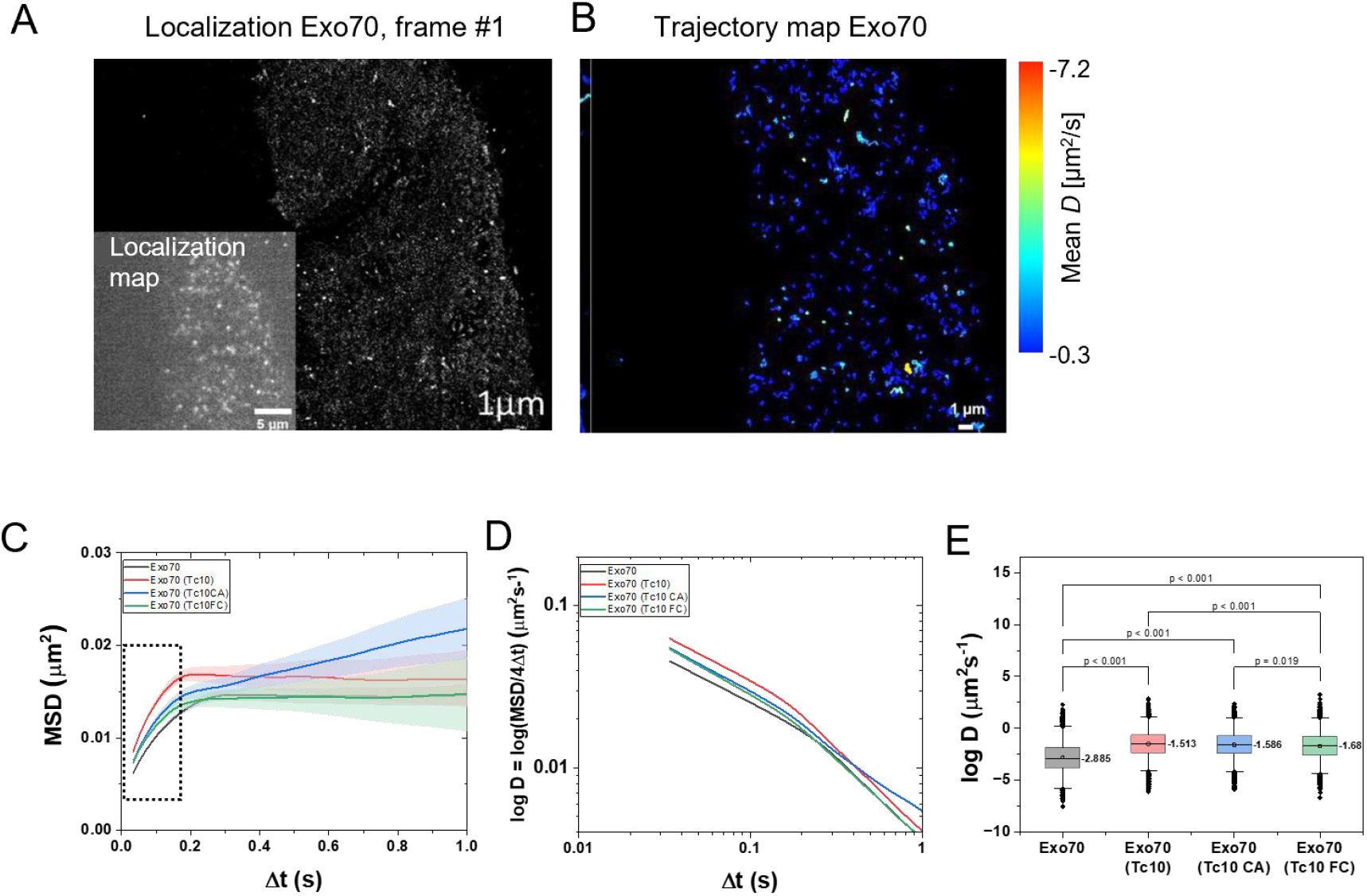
Dynamics of Exo70 in the presence of TC10 and its mutants in HeLa cells. (A) Localization of Exo70 in HeLa cells. One frame showing single Exo70 particles. Inset: Cumulative Localization map of Exo70, 5000 frames, frame rate 17 ms, pixel size: 65 nm. (B) Trajectory map of Exo70 (85 s). (C) Ensemble time-averaged MSD curves for Exo70 (black curve), Exo70 with TC10 (red curve), Exo70 with TC10CA (blue curve), and Exo70 with TC10 FC (green curve). The shaded area in the corresponding colour represents the standard error of the mean. (D) Changes of diffusion coefficients (logarithmic) over time. (E) Comparison of the Exo70 diffusion in the presence of different TC10 variants. The median values of the diffusion coefficients are presented. The box limit indicates the 25th and 75th percentiles; whiskers extend to 1.5 times the interquartile range of the 25th and 75th percentiles; a straight line in the box plot gives the median values and this value is presented next to the box; the significant p values were estimated using the Kruskal-Wallis ANOVA test followed by post-hoc analysis. N=5.

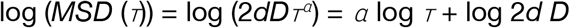

, where *D* is the diffusion coefficient, *⊤* is the time and *a* is the anomalous coefficient [12]. Only the first 200 ms of the trajectories were fitted and only those fits were selected for which the quality of the fit, R^2^ coefficients were > 0.8. These fits provided the values of *D* for individual tracks according to the models described by Qian **[13]** (**Figure 1E**) (**Table S1**). In Hela cells, TC10 and TC10 mutants increased the mobility of Exo70 at the PM, whereby TC10CA had the strongest effect.

### TC10 decreases Exo70 mobility in the axonal growth cone of cortical neurons

Next, we determined the dynamics of Exo70 in the axonal growth cone of cortical neurons. Neurons were prepared from the cortical region of rat embryos at day 18 and co-transfected with Exo70-HaloTag and Snap-TC10, respectively TC10 variants, at DIV1. SPT was conducted at DIV2 in the growth cone which is the site of membrane expansion. We only analyzed neurons with one axon. Single particles of Exo70 were recorded in the absence and presence of TC10, TC10CA, and TC10FC, respectively (**Figure 2A**). Movies of 5000 frames were recorded (58.8 Hz acquisition) and particles were localized. The superimposition of all Exo70 particles resulted in a localization map (**Figure 2B**). From the data, trajectories were generated. Individual trajectories were color-coded based on the mean velocity of the particle. All trajectories of one movie were superimposed to generate a trajectory map. The pattern of the localization and trajectory maps suggest that Exo70 may be localized along actin filaments (**Figure 2C**). We determined the mean square displacements (MSD) (**Figure 2D**) and plotted MSD vs. time on a logarithmic scale (**Figure 2E**) to show the deviation from normal behavior.

**Figure 2:**
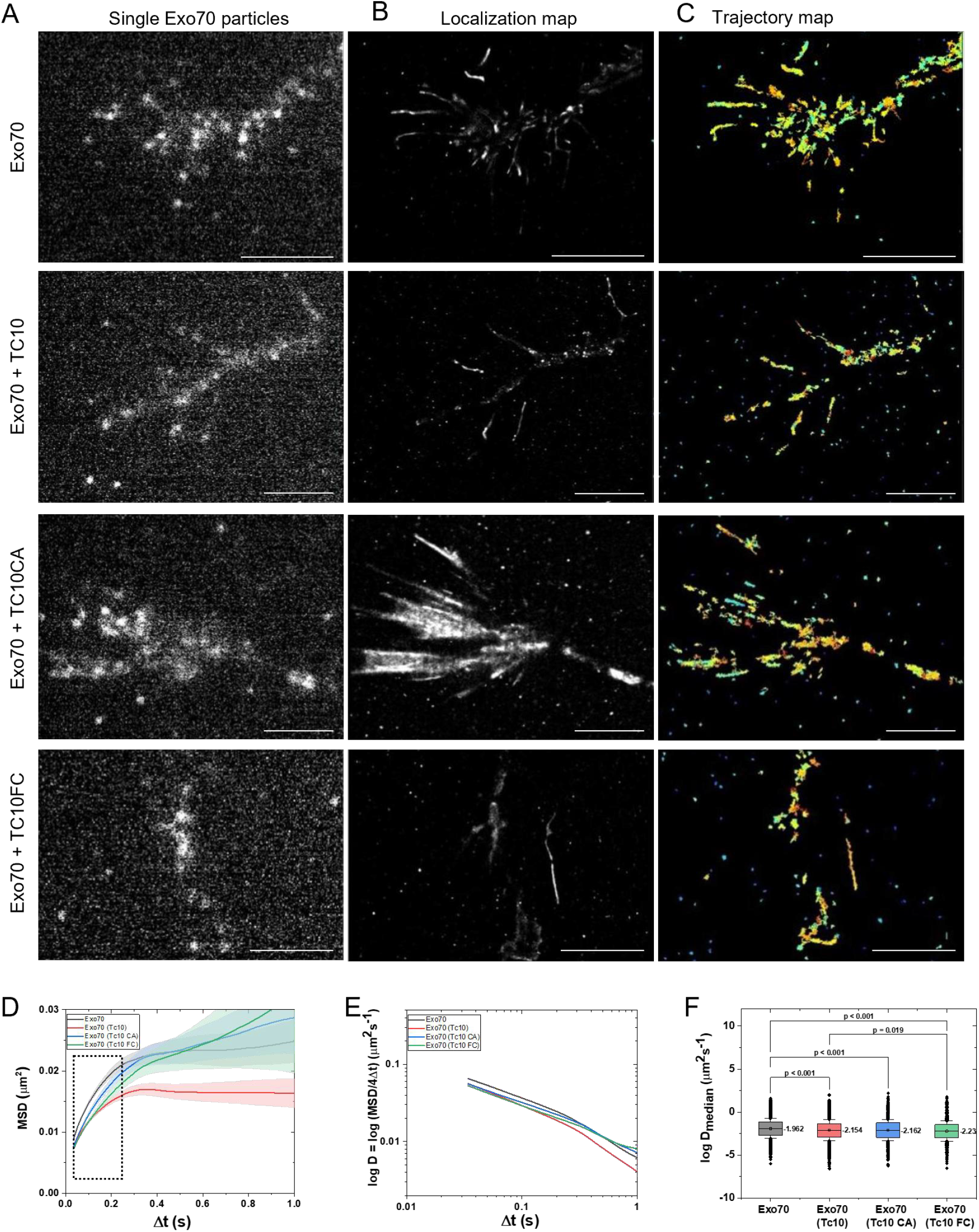
Effect of TC10 on Exo70 dynamics in the axonal growth cone of cortical neurons. (A) Single particle imaging of Exo70 in the absence and presence of TC10 and TC10 variants in the growth cone, a single frame is shown. (B) Cumulative image of 5000 frames with localized Exo70 particles. (C) Trajectory map of Exo70 in the absence and presence of TC10 and TC10 variants. Each trajectory is labelled in a colour related to the mean diffusion coefficient. Cumulative image of 5000 frames recoded with 58.8 Hz. (D) Characterization of Exo70 diffusion in the growth cone: mean square displacements (MSD) over time. (E) Change of the diffusion coefficient D over time (-log D). (F) Diffusion coefficients as -log D. N=2.

Exo70 mobility was significantly decreased in the presence of TC10 (**Table S2**). TC10CA and TC10FC had similar effects on Exo70 dynamics in the growth cones of cortical axons at DIV2 (**Figure 2D**). Diffusion coefficients of Exo70 were significantly lower in the presence of TC10 and active TC10 variants (**Table S2**). This indicates an immobilization of Exo70 due to vesicle tethering at the PM, promoted by TC10 and its active derivatives **[6, 7]**. Also, in the soma of the cortical neurons, TC10 overexpression reduced Exo70 mobility at the plasma membrane (Figure S2). However, the radius of confinement of Exo70 differed, reflecting that Exo70 occupies different plasma membrane domains in the soma and growth cone.

### TC10 has no impact on Exo70 dynamics in growth cones of hippocampal neurons

To determine the effect of TC10 on Exo70 dynamics in the growth cone of hippocampal neurons, hippocampal neurons from E18 rat embryos were transfected with Exo70-HaloTag without or with SNAP-TC10 and its variants and Exo70 dynamics was determined as before by SPT. In the case of hippocampal neurons, Exo70 dynamics in the growth cone was not affected by TC10 (**Figure 3, Table S3**), suggesting that TC10 plays a minor role for Exo70 dynamics.

**Figure 3:**
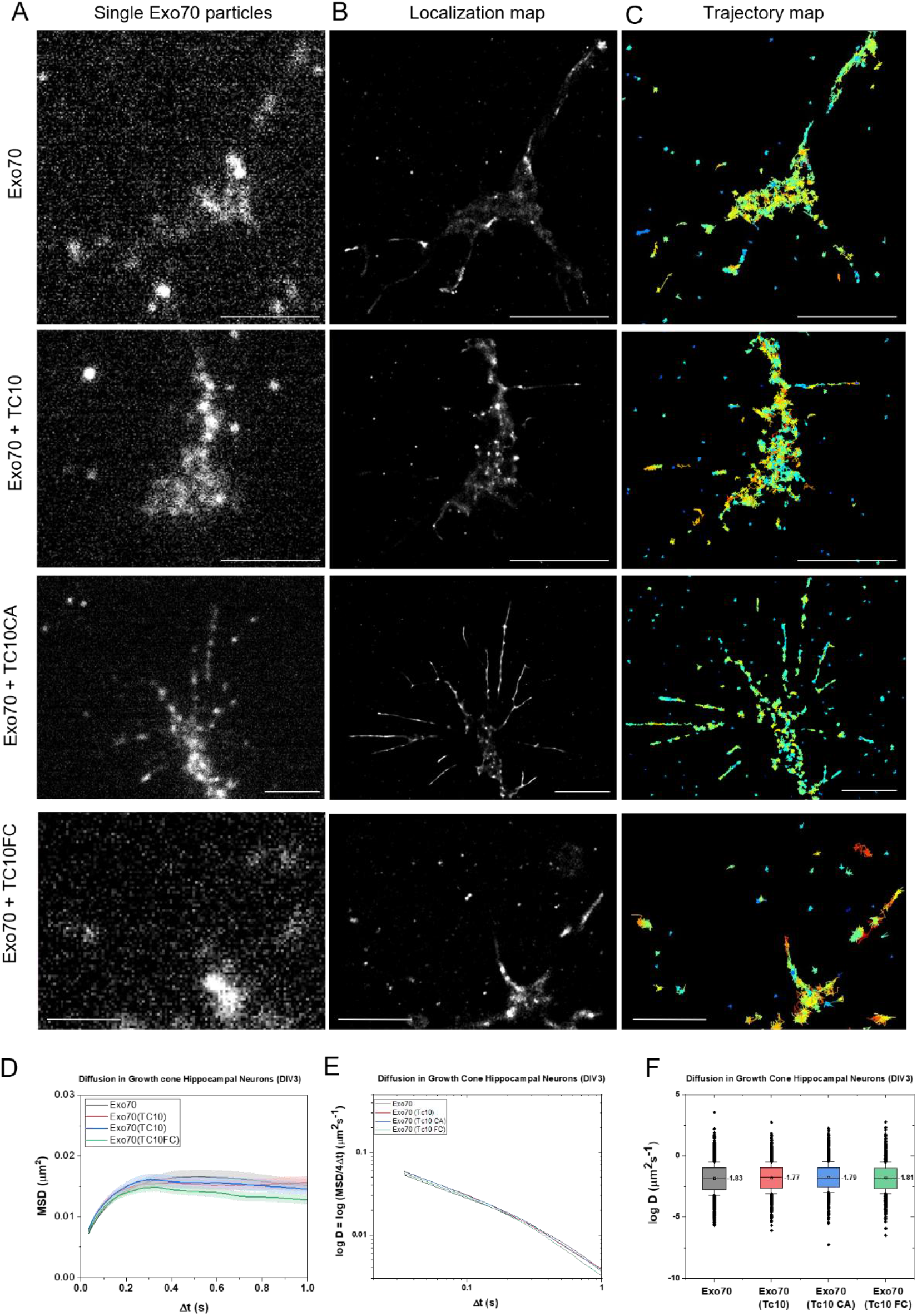
Effect of TC10 on Exo70 dynamics in the axonal growth cone of hippocampal neurons. (A) Single particle imaging of Exo70 in the absence and presence of TC10 and TC10 variants in the growth cone, single frame shown. (B) Cumulative image of 5000 frames with localized Exo70 particles. (C) Trajectory map of Exo70 in the absence and presence of TC10 and TC10 variants. Each trajectory is labelled in a colour related to the mean diffusion coefficient. Cumulative image of 5000 frames recoded with 58.8 Hz. (D) Characterization of Exo70 diffusion in the growth cone: mean square displacements (MSD) over time. (E) Change of the diffusion coefficient D over time (-log D). (F) Diffusion coefficients as -log D. N=2. Scale bars: 5 µm (A-C).

### TC10 increases the fraction of Exo70 particles showing anomalous motion in cortical neurons

The log*D*/Dt plots of Exo70 diffusion (**Figure 1E, 2E, 3E**) showed non-linear behaviour indicating anomalous diffusion. Anomalous diffusion can be caused by obstacles, structural constraints of the microcompartment, or by interactions that affect diffusion behavior, such as increased crowding **[14]**. To dissect that further, we determined the alpha coefficients (a) from MSD curves for all conditions (**Figure 4**). a is a measure of the type of movement, and it is expected that Exo70/exocyst tethering is associated with anomalous diffusion. a coefficients were lowest for Exo70 diffusion in the cortical growth cone. Under the influence of TC10FC, the a coefficient for Exo70 diffusion further decreased in cortical axon growth cones (**Figure 4A**). TC10 had not effect on a of Exo70 motion in hippocampal neurons. The fraction of Exo70 particles displaying anomalous motion (a<1) was highest in cortical axons (**Figure 4B**). in line with no change in *D* in the presence of TC10 and its functional variants (**Figure 3**). The anomalous diffusing Exo70 particles decreased in HeLa cells in the presence of TC10 from 72 % to 60 %, in the presence of TC10CA to 61 % and in the presence of TC10FC to 62 %. Contrary, in cortical neurons, the fraction of anomalous diffusing Exo70 particles increased from 74 % (no TC10 OE), to 77 % with TC10, 76 % (with TC10CA) and 80 % with TC10FC (**Figure 4B**). No change in anomalous diffusion was observed in hippocampal axon growth cones. Together, this data confirms that TC10 affects Exo70 dynamics in axonal growth cones of cortical neurons but not in hippocampal neurons.

**Figure 4:**
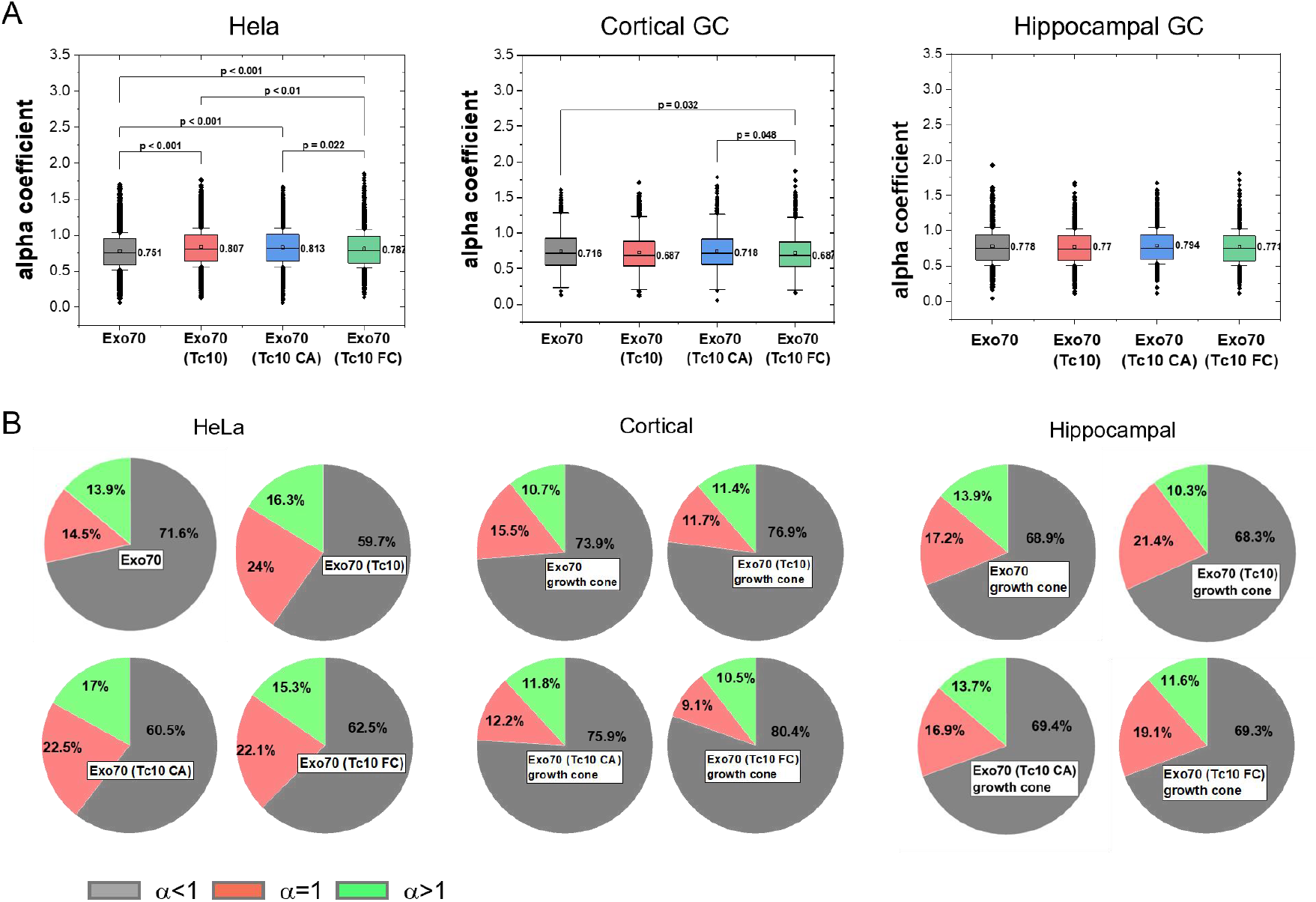
Anomalous diffusion of Exo70 at the plasma membrane. (A) Mean alpha coefficients (a) for Exo70 in HeLa cell and the growth cone of cortical, respectively hippocampal neurons, in the absence and presence of TC10 and its variants. Box-plots present the mean values of the alpha coefficient and the box limit indicates the 25th and 75th percentiles; whiskers extend to 1.5 times the interquartile range of the 25th and 75th percentiles; a square in the box plot gives the mean values and this value is presented next to the box; the significant p values are estimated using one ANOVA test followed by Tukey’s test analysis. Replicates: N=5 (HeLa), N=2 (cortical neurons), N=3 (hippocampal neurons). (B) Dissection of motion behavior. The pie chart shows the respective population of Exo70 following anomalous or confined motion (grey part, a<1), Brownian motion (red part, a=1) and directed motion (green portion, a>1).

## Discussion

In this study, we compared the impact of the regulatory Rho-GTPase TC0 on the dynamics of Exo70 at the plasma membrane in non-polarized cells and in the axonal growth cone of cortical and hippocampal neurons during polarization. The axonal growth cone is a zone of membrane expansion that requires the interaction of the activating Rho-GTPase TC10 with the exocyst subunit Exo70 **[6]**. In HeLa cells, TC10 increased the diffusion of Exo70. Since our HeLa cells were not EGF-stimulated **[5]**, the TC10 effect on Exo70 likely concern a population that is not incorporated into the exocyst complex and has function beyond exocytosis **[15]**. For cortical neurons, we found that TC10 decreased the diffusion of Exo70 in the soma and in the growth zone. Moreover, the fraction of Exo70 that displayed anomalous diffusion (a<1) was increased. The reduced mobility of Exo70 linked to TC10 expression can be explained by enhanced Exo70/exocyst vesicle tethering. In this scenario, GTP-TC10 interacts with Exo70 to promote vesicle binding. The GTP-TC10 variant TC10-Q67L, which is constitutively active (TC10CA), had the same effect as TC10 on Exo70 mobility. However, the fast-cycling variant TC10/34L increased the fraction of Exo70 displaying confined motion in accordance with promotion of vesicle fusion **[7]**. The confinement radius of Exo70 was larger in n the growth cone than in the soma or in Hela cells (Figure S3). This is likely determined by the interaction of Exo70 with the cytoskeleton, which forms extended patterns in the growth cone **[16, 17]**. In contrast to the growth cone of cortical neurons, overexpression of TC10 had no effect on Exo70 dynamics in the growth cone of hippocampal neurons. Although a previous study reported TC10 control of Exo70 in hippocampal neurons in **[6]**, axonal growth is also regulated by Cdc42b **[8]**. Apparently, the integrated action of both Rho-family GTPases is specifically crucial in axonal outgrowth of hippocampal neurons **[18, 19]**.

## Availability of data and materials

All data supporting the conclusions of this article are included within the manuscript and raw data and movies are available upon reasonable request to the corresponding author.

## Appendices

Supplementary information is found online.

## Author contributions

HG and KB designed experiments – HG conducted experiments – KB and HG analyzed data – KB and HG wrote the manuscript.

## Acknowledgements

The authors were members of the CIM. KB and HG were funded by the SFB1348. HG was supported by the GRC grant #386797833. We are thankful to Priyadarshini Ravindran and Andreas Püschel for providing transfected neurons and the Exo70 and TC10 mutants.

## References

1. Barnes, A.P. and F. Polleux, Establishment of axon-dendrite polarity in developing neurons. Annu Rev Neurosci, 2009. 32: p. 347–81.

2. Dotti, C.G., C.A. Sullivan, and G.A. Banker, The establishment of polarity by hippocampal neurons in culture. J Neurosci, 1988. 8(4): p. 1454–68.

3. Rakic, P., et al., Computer-aided three-dimensional reconstruction and quantitative analysis of cells from serial electron microscopic montages of foetal monkey brain. Nature, 1974. 250(461): p. 31–4.

4. Affenzeller, M.J., et al., Salt stress-induced cell death in the unicellular green alga Micrasterias denticulata. J Exp Bot, 2009. 60(3): p. 939–54.

5. Kawase, K., et al., GTP hydrolysis by the Rho family GTPase TC10 promotes exocytic vesicle fusion. Dev Cell, 2006. 11(3): p. 411–21.

6. Dupraz, S., et al., The TC10-Exo70 complex is essential for membrane expansion and axonal specification in developing neurons. J Neurosci, 2009. 29(42): p. 13292–301.

7. Fujita, A., et al., GTP hydrolysis of TC10 promotes neurite outgrowth through exocytic fusion of Rab11- and L1-containing vesicles by releasing exocyst component Exo70. PLoS One, 2013. 8(11): p. e79689.

8. Ravindran, P. and A.W. Puschel, An isoform-specific function of Cdc42 in regulating mammalian Exo70 during axon formation. Life Sci Alliance, 2023. 6(3).

9. Liss, V., et al., Self-labelling enzymes as universal tags for fluorescence microscopy, super-resolution microscopy and electron microscopy. Sci Rep, 2015. 5: p. 17740.

10. Inoue, M., et al., The exocyst complex is required for targeting of Glut4 to the plasma membrane by insulin. Nature, 2003. 422(6932): p. 629–33.

11. Lopez Tobon, A., et al., The guanine nucleotide exchange factor Arhgef7/betaPix promotes axon formation upstream of TC10. Sci Rep, 2018. 8(1): p. 8811.

12. Saxton, M.J., A biological interpretation of transient anomalous subdiffusion. II. Reaction kinetics. Biophys J, 2008. 94(3): p. 760–71.

13. Qian, H., M.P. Sheetz, and E.L. Elson, Single particle tracking. Analysis of diffusion and flow in two-dimensional systems. Biophys J, 1991. 60(4): p. 910–21.

14. Guigas, G. and M. Weiss, Influence of hydrophobic mismatching on membrane protein diffusion. Biophys J, 2008. 95(3): p. L25-7.

15. Zhu, Y., B. Wu, and W. Guo, The role of Exo70 in exocytosis and beyond. Small GTPases, 2019. 10(5): p. 331–335.

16. Fritzsche, M., et al., Self-organizing actin patterns shape membrane architecture but not cell mechanics. Nat Commun, 2017. 8: p. 14347.

17. Nozumi, M., et al., Coordinated Movement of Vesicles and Actin Bundles during Nerve Growth Revealed by Superresolution Microscopy. Cell Rep, 2017. 18(9): p. 2203–2216.

18. Gracias, N.G., N.J. Shirkey-Son, and U. Hengst, Local translation of TC10 is required for membrane expansion during axon outgrowth. Nat Commun, 2014. 5: p. 3506.

19. Erschbamer, M.K., C.P. Hofstetter, and L. Olson, RhoA, RhoB, RhoC, Rac1, Cdc42, and Tc10 mRNA levels in spinal cord, sensory ganglia, and corticospinal tract neurons and long-lasting specific changes following spinal cord injury. J Comp Neurol, 2005. 484(2): p. 224–33.

