## supporting material for "TC10 differently controls the dynamics of Exo70 in growth cones of cortical and hippocampal neurons"

#### Supplementary Figures

**Figure S 1: Colocalization of Exo70 and TC10 in HeLa cells**. (a) Endogenous TC10 (green) and (b) endogenous Exo70 (red), immune-staining. (c) Merge of Exo70 and TC10. TC10 and Exo70 are found both at the plasma membrane and as puncta. (d, e, f) Zoomed in regions of figures (a, b, c) (white boxes), size 100x100 pixels. Colocalization of Exo70 and TC10 are shown as yellow spots in (c) and (f). Primary/secondary antibody are Exo70/AF555, TC10/AF488. (g) The threshold for both the green (TC10) and red (Exo70) channels was adjusted, and colocalization was performed in Imaris™ (Bitplane) in an area of 100x100 pixels. Colocalization is indicated by white spots. (h) Scatter plot to determine the Pearson's coefficient. (i) Calculated colocalization (%) averaged over 30 images (100x100 px). Scale bars: 10 µm (a,b,c), 1 µm (d,e,f).

**Figure S 2**: **Effect of TC10 on Exo70 dynamics in the soma of cortical neurons**. (A) Single particle imaging of Exo70 in the absence and presence of TC10 and TC10 variants in the soma, a single frame is shown. (B) Cumulative image of 5000 frames with localized Exo70 particles. (C) Trajectory map of Exo70 in the absence and presence of TC10 and TC10 variants. Each trajectory is labelled in a colour related to the mean diffusion coefficient. Cumulative image of 5000 frames recoded with 58.8 Hz. (D) Characterization of Exo70 diffusion in the soma: mean square displacements (MSD) over time. (E) Change of the diffusion coefficient *D* over time (-log D). (F) Diffusion coefficients as -log *D*. The box limit indicates the 25th and 75th percentiles; whiskers extend to 1.5 times the interquartile range of the 25th and 75th percentiles; a straight line in the box plot gives the median values and this value is presented next to the box; the significant p values were estimated using the Kruskal-Wallis ANOVA test followed by post-hoc analysis. N=2.

**Figure S 3: Confinement and anomalous diffusion of Exo70 at the plasma membrane in HeLa cells and cortical neurons.** (A) Means square displacements. (B) Dissection of motion behavior. The pie chart shows the respective population of Exo70 following anomalous or confined motion (grey part, $\alpha<1$), Brownian motion (red part, $\alpha=1$) and directed motion (green portion, $\alpha>1$).

**Figure S 1**

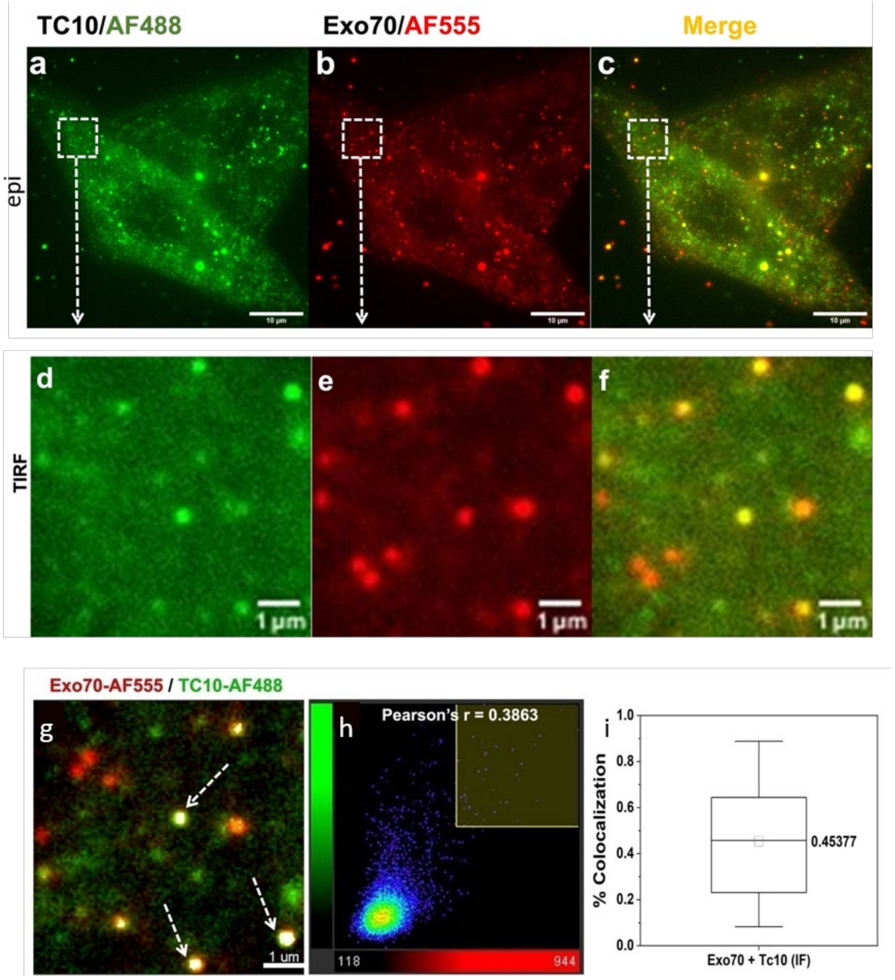
TC10/AF488
Exo70/AF555
Merge
a
b
c
epi
d
e
f
TIRF
1 µm
1 µm
1 µm
Exo70-AF555 / TC10-AF488
g
1 um
h
Pearson's r = 0.3863
118
944
i
% Colocalization
1.0
0.8
0.6
0.4
0.2
0.0
0.45377
Exo70 + Tc10 (IF)

### Figure S 2

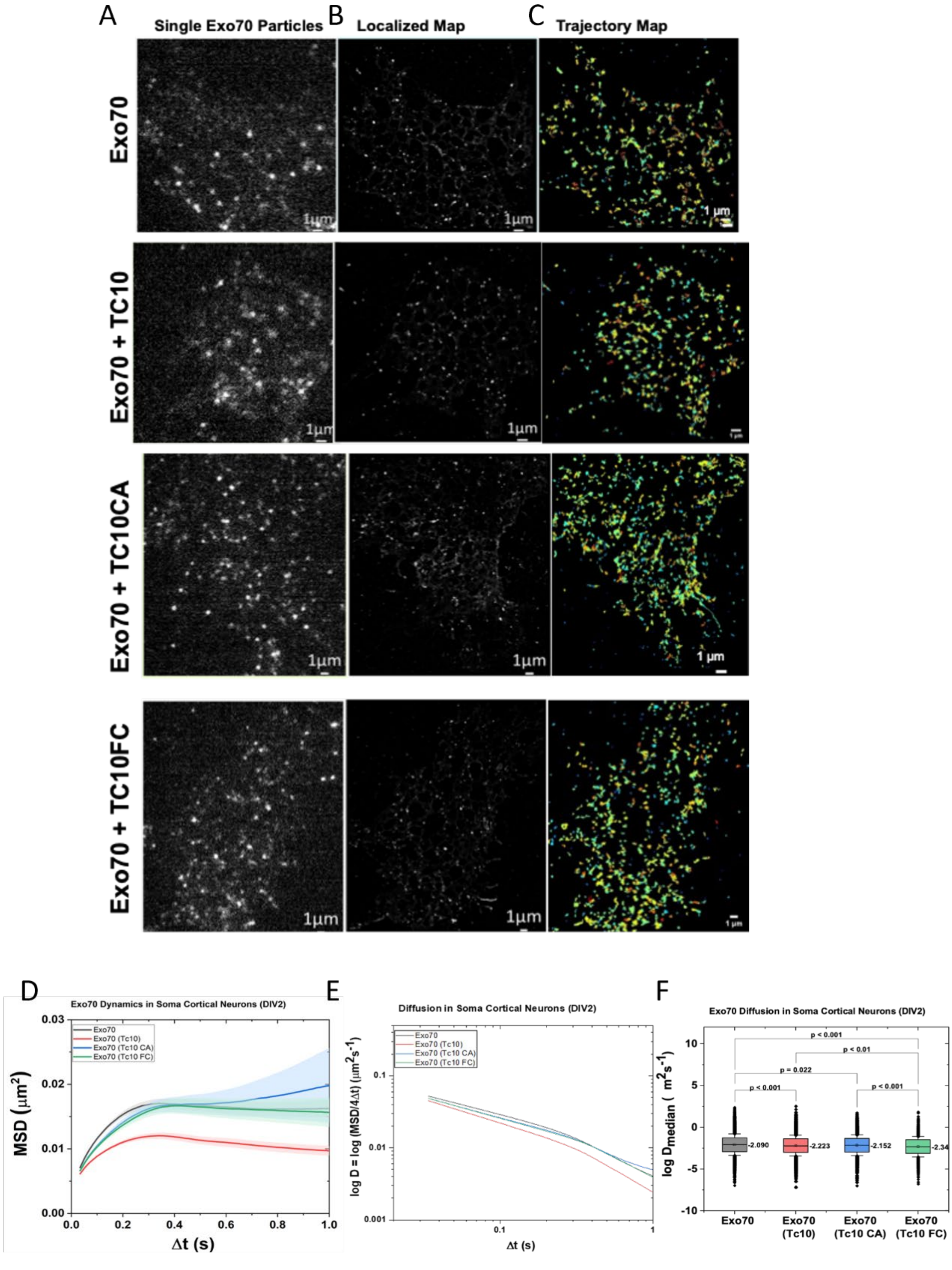
A
Single Exo70 Particles
B
Localized Map
C
Trajectory Map
Exo70
Exo70 + TC10
Exo70 + TC10CA
Exo70 + TC10FC
1μm
D
Exo70 Dynamics in Soma Cortical Neurons (DIV2)
MSD (μm²)
Δt (s)
Exo70
Exo70 (Tc10)
Exo70 (Tc10 CA)
Exo70 (Tc10 FC)
E
Diffusion in Soma Cortical Neurons (DIV2)
log D = log (MSD/4Δt) (μm²s⁻¹)
Δt (s)
F
Exo70 Diffusion in Soma Cortical Neurons (DIV2)
log Dmedian ( m²s⁻¹)
p < 0.001
p < 0.01
p = 0.022
p < 0.001
p < 0.001
-2.090
-2.223
-2.152
-2.34
Exo70
Exo70 (Tc10)
Exo70 (Tc10 CA)
Exo70 (Tc10 FC)

### Figure S 3

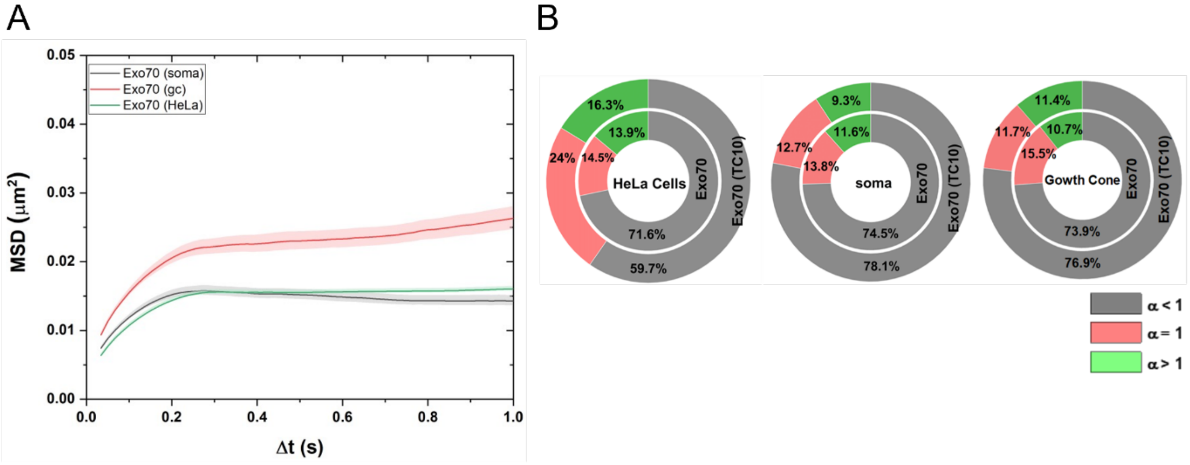
A
B
Exo70 (soma)
Exo70 (gc)
Exo70 (HeLa)
MSD (μm²)
Δt (s)
0.05
0.04
0.03
0.02
0.01
0.00
0.0
0.2
0.4
0.6
0.8
1.0
16.3%
13.9%
24%
14.5%
HeLa Cells
Exo70
Exo70 (TC10)
71.6%
59.7%
9.3%
11.6%
12.7%
13.8%
soma
74.5%
78.1%
11.4%
10.7%
11.7%
15.5%
Gowth Cone
73.9%
76.9%
α < 1
α = 1
α > 1

*Table 1: Effect of TC10 on the diffusion of Exo70 in nonpolar HeLa cells, N=3.*

| HeLa cells | Total # tracks | $\mathcal{D}_{median}$±s.e.m. (μm²s⁻¹) | $\alpha_{mean}$ ± s.e.m. | Number of cells |
|---|---|---|---|---|
| Exo70 | 22724 | 0.011 ± 0.001 | 0.776 ± 0.003 | 20 |
| Exo70 (TC10) | 11644 | 0.031 ± 0.002 | 0.832 ± 0.005 | 20 |
| Exo70 (TC10CA) | 9249 | 0.026 ± 0.003 | 0.830 ± 0.005 | 20 |
| Exo70 (TC10FC) | 16188 | 0.021 ± 0.002 | 0.811 ± 0.004 | 20 |

*Table 2: Effect of TC10 on the diffusion of Exo70 in the axonal growth cone of cortical neurons, N=2.*

| Growth Cone Cortical Neurons (DIV2) | Total tracks | $\mathcal{D}_{median}$±s.e.m. ($\mu m^2 s^{-1}$) | $\alpha_{mean}$ ± s.e.m. | Number of cells |
|---|---|---|---|---|
| **Exo70** | 6938 | 0.011 ± 0.002 | 0.749 ± 0.006 | 13 |
| **Exo70 (TC10)** | 9108 | 0.007 ± 0.003 | 0.731 ± 0.006 | 13 |
| **Exo70 (TC10CA)** | 4402 | 0.007 ± 0.005 | 0.745 ± 0.007 | 12 |
| **Exo70 (TC10FC)** | 4949 | 0.006 ± 0.003 | 0.724 ± 0.007 | 12 |

*Table 3: Effect of TC10 on the diffusion of Exo70 in the axonal growth cone of hippocampal neurons. N=2.*

| Growth Cone Hippocampal Neurons (DIV3) | Total tracks | $\mathcal{D}_{median}$±s.e.m. ($\mu m^2 s^{-1}$) | $\alpha_{mean}$ ± s.e.m. | Number of cells |
|---|---|---|---|---|
| **Exo70** | 5544 | 0.015 ± 0.001 | 0.778 ± 0.007 | 9 |
| **Exo70 (TC10)** | 3407 | 0.017 ± 0.002 | 0.770 ± 0.009 | 8 |
| **Exo70 (TC10CA)** | 5229 | 0.016 ± 0.003 | 0.794 ± 0.008 | 8 |
| **Exo70 (TC10FC)** | 4552 | 0.016 ± 0.003 | 0.771 ± 0.008 | 8 |
